# Correcting index databases improves metagenomic studies

**DOI:** 10.1101/712166

**Authors:** Guillaume Méric, Ryan R. Wick, Stephen C. Watts, Kathryn E. Holt, Michael Inouye

**Author notes:** These authors have contributed equally (Guillaume Méric, Ryan R. Wick).

## Abstract

Assessing the taxonomic composition of metagenomic samples is an important first step in understanding the biology and ecology of microbial communities in complex environments. Despite a wealth of algorithms and tools for metagenomic classification, relatively little effort has been put into the critical task of improving the quality of reference indices to which metagenomic reads are assigned. Here, we inferred the taxonomic composition of 404 publicly available metagenomes from human, marine and soil environments, using custom index databases modified according to two factors: the number of reference genomes used to build the databases, and the monophyletic strictness of species definitions. Index databases built following the NCBI taxonomic system were also compared to others using Genome Taxonomy Database (GTDB) taxonomic redefinitions. We observed a considerable increase in the rate of read classification using modified reference index databases as compared to a default NCBI RefSeq database, with up to a 4.4-, 6.4- and 2.2-fold increase in classified reads per sample for human, marine and soil metagenomes, respectively. Importantly, targeted correction for 70 common human pathogens and bacterial genera in the index database increased their specific detection levels in human metagenomes. We also show the choice of index database can influence downstream diversity and distance estimates for microbiome data. Overall, the study shows a large amount of accessible information in metagenomes remains unexploited using current methods, and that the same data analysed using different index databases could potentially lead to different conclusions. These results have implications for the power and design of individual microbiome studies, and for comparison and meta-analysis of microbiome datasets.

## Introduction

For more than 3 billion years, microbes have established complex ecological niches in environments and hosts throughout the planet. This makes them ubiquitous components of biogeochemical cycles on land [1], in the sea [2], the atmosphere [3], and on or inside other living organisms [4, 5] including humans, in which they are important for development and health [6, 7]. However, technical constraints limit our ability to study the ecology of microorganisms, in particular the widespread lack of suitable culturing methods [8]. An important advance in the analysis of microbial communities has been the use of sequence-based, culture-independent methods to study the diversity and composition of clinical and environmental samples and their biological functions. The increasing affordability of high-throughput sequencing has led to an increase in metagenomics studies, in which a sample’s total extracted DNA can be sequenced as a whole. Accurately determining and quantifying the taxonomic composition of a metagenome is a critical first step in many analyses, such as the association with host phenotype, host genotype, disease status or environmental properties.

Metagenomic classification begins with the accurate assignment of sequencing reads to a reference database, or “index”, comprising reference genomes and their corresponding taxonomic definitions. A wealth of metagenomic classification algorithms have been developed in the last few years [4, 9–16], mainly focusing on improving classification speed and memory usage, including popular methods such as Kraken [17, 18] or Centrifuge [19]. A given read sequence may be shared among closely-related species, particularly when the read length is short, and so classifiers can assign reads to the last common ancestor (LCA) of all taxa sharing their sequence (“LCA-classification”). Despite the development of ever-more efficient classifying algorithms and tools, comparatively little has been done to improve the quality of the reference indices used to define the taxa to which reads are assigned. Recent efforts showed that the addition of new genomes to NCBI RefSeq could influence metagenomic classification performance, with indices built on most recent releases of NCBI RefSeq able to classify more reads overall, but fewer at the species level [20]. Generally, most methods and studies use a selection of representative, often complete genomes from curated repositories to build indices from all described bacteria and archaea using their reported taxonomic definitions [16, 21], typically NCBI RefSeq [22] for whole representative genomes, and SILVA, Greengenes, or RDP [23] for 16S rRNA-based studies.

Defining accurate monophyletic bacterial species boundaries has always been a challenge. Bacterial taxonomy has historically been defined using imprecise biochemical or ecological phenotypes, with more recent genotyping studies offering numerous examples of clustered “species” previously thought to be distinct, and *vice versa* [24–27]. As a result, microbial taxonomies in reference repositories are riddled with inconsistencies, with described taxa often forming polyphyletic groupings [28, 29], necessitating reconciliation between microbial systematics and genomics [30]. This has recently been addressed by redefining taxonomic definitions using a phylogenetic depth coefficient inferred from a robust prokaryotic phylogeny [28]. This effort, summarised in the Genome Taxonomy Database (GTDB), aims to define strictly monophyletic species groups of equivalent phylogenetic depth. It produced a wealth of novel definitions at various taxonomic levels of the microbial tree of life, redefining approximately 58% of all previous NCBI-based taxonomic definitions [28].

Most classification tools will recommend the use of default indices, built using a set of complete genomes from NCBI RefSeq. In this study, we assessed the potential for improvement and addressed the following questions: does the choice of reference index affect the performance of metagenomic classification? Does the addition of draft reference genome sequences improve classification? Should we use default NCBI-based indices, custom human microbiome-enhanced indices; or GTDB-based indices? Is the inclusion of metagenome-assembled genomes (MAGs) beneficial? Is the strict monophyly of taxonomic definitions in indices important for classification performance? To answer these questions, we created seven custom indices (**Table 1**) using NCBI-based and GTDB-based taxonomic definitions, and examined their classification performance on samples from three diverse and representative metagenomic datasets: human body sites, marine and soil. Our work addresses the metagenomic classification bias, whereby sequencing reads for particular taxa are present in metagenomics data but remain unclassified using current methods and recommendations. This has important consequences for the classification of metagenomic datasets and downstream applications such as microbiome-wide association studies.

**Table 1.** Description of the seven classification indices used in this study. The release numbers correspond to NCBI RefSeq releases of genomes from which the reference genomes used to build indices were obtained.

| Index name | Reference database and taxonomic definitions used | Description | Total number of reference genomes included | Strictly monophyletic species definitions |
| --- | --- | --- | --- | --- |
| NCBI_r86 | NCBI RefSeq release 86 | Complete microbial genomes (r86) | 8674 | N |
| GTDB_r86_8.6k | GTDB release 86 | Same genomes as NCBI_8.6k but with GTDB taxonomic definitions, to compare effect of strict monophyletic definitions | 8674 | Y |
| NCBI_r88 | NCBI RefSeq release 88 | Complete microbial genomes (r88) | 10089 | N |
| NCBI_r88_Human17k | NCBI RefSeq release 88 | Same as NCBI_r88 with the addition of all draft genomes from 70 genera of interest, strictly corrected for monophyly | 16908 | Y (for 70 genera only) |
| GTDB_r86_noMAGs | GTDB release 86 | GTDB r86 without metagenome-assembled genomes (MAGs) | 25660 | Y |
| GTDB_r86 | GTDB release 86 | All dereplicated* bacterial and archaeal genomes used to curate the GTDB taxonomy in the GTDB study | 28560 | Y |
| GTDB_r86_46k | GTDB release 86 | Manual dereplication* of GTDB release 86 to get more bacterial and archaeal reference genomes with GTDB taxonomic definitions | 46006 | Y |

\* Dereplication is defined as the threshold-based selection of representative reference genomes for phylogenetically-similar clusters.

## Results

### Substantial improvements in classification performance can be achieved using larger indices

To examine the impact of custom indices on metagenomic classification performance, we classified 404 metagenomic samples from three different datasets using seven custom indices (**Figure S1, Table 1**) and quantified the proportion of reads per sample that were classified to any taxon and the proportion that remained unclassified (**Figure 1, Figure S2**). Our custom index databases were corrected for two distinct factors: (a) number of reference genomes for each species used to build the index, and (b) strict monophyletic species definition for these reference genomes (**Table 1**). We observed a drastic improvement in classification performance using custom indices, built with more reference genomes, i.e. the greater the number of reference genomes used to build the index, the greater the proportion of reads classified (**Figure 1A-C**). This effect was not associated with sequencing depth (**Figure S3**). For instance, using the NCBI_r88_Human17k index on human metagenomes, which includes only 1.67-fold more genomes than NCBI_r88 (selectively chosen from 70 known human microbiome taxa) and monophyly correction (**Table 1**), the median proportion of classified reads per sample increased from 54.7% to 76.5%. The index built with the largest number of genomes, GTDB_r86_46k, consistently classified the most reads in every sample tested. The increase in the median percentage of classified reads per sample from a default NCBI_r86 classification for human metagenomes was from 53.6% to 91.3% (median increase of +69.4%; range of +3.9% to +342.8%) (**Figure S2A, Table S2, Table S3**). Similarly, the increase in classified reads per sample for marine metagenomes was from a median of 14.1% to a median of 55.2% (median increase of +276.2%; range of +94.6% to +536.3%); and in soil metagenomes from 33.2% to 66.3% (median increase of +100.7% reads/sample; range of +85.7% to +120.6%) (**Figure S2B-C, Table S2, Table S3**).

**Figure 1.**
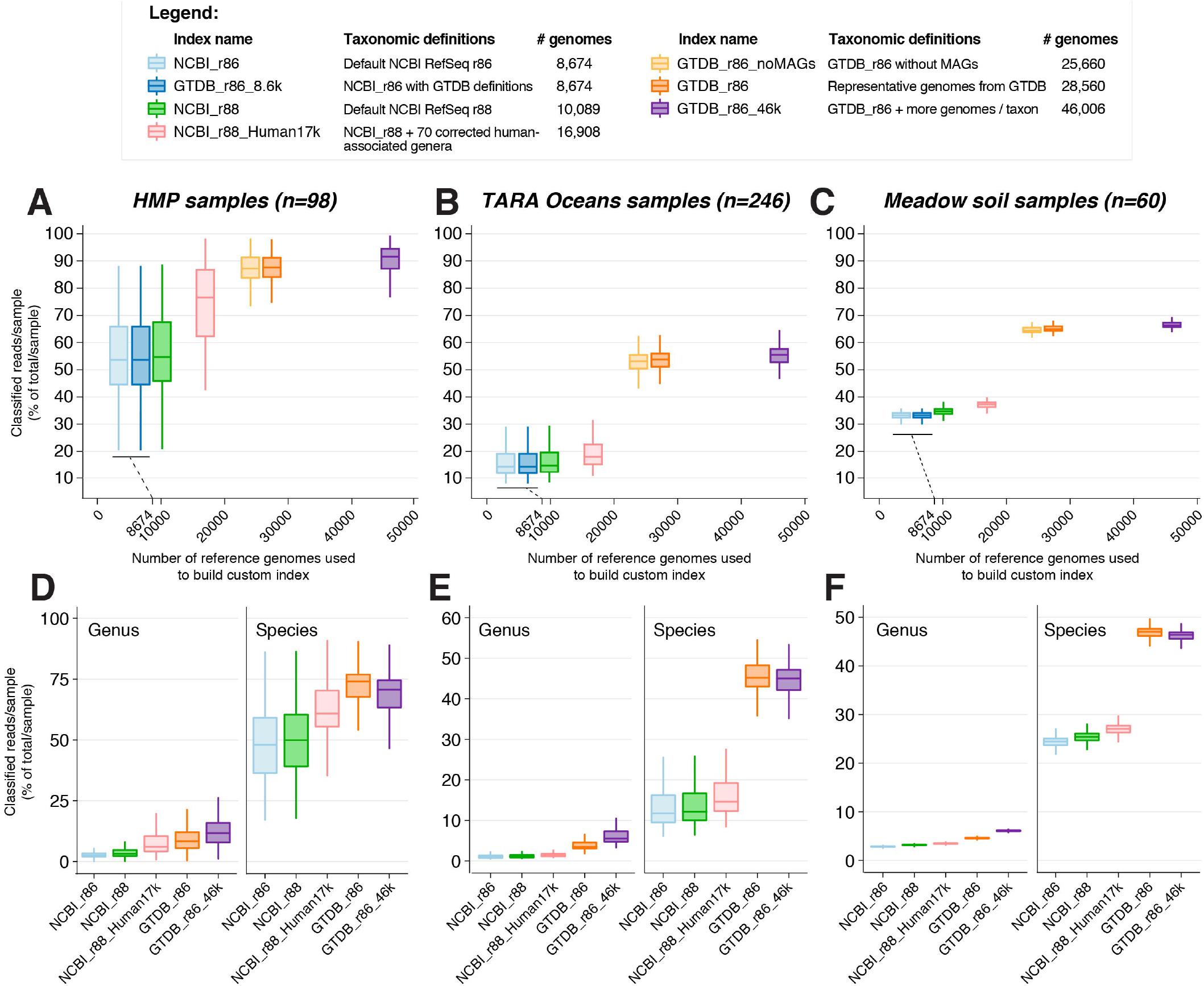
Large index databases substantially improve metagenomic classification performance and accuracy, including at lower taxonomic levels. Sequencing reads from three datasets (HMP samples, n=98; TARA Oceans samples, n=246; meadow soil samples, n=60) were classified using the seven index databases presented in Table 1. Boxplots (Tukey) show the distribution of the proportion of unclassified and classified reads/samples for human samples (A), marine samples (B) and soil samples (C) using seven indices (y-axis) is shown for each index size (x-axis), defined as the number of reference genomes used to build the index. Distributions of the breakdown of read classification to the two lowest taxonomic levels (genus, species) for human samples (D), marine samples (E) and soil samples (F) are shown for the NCBI_r86 default index (light blue), two indices based on NCBI_r88 (NCBI_r88 in green and NCBI_r88_Human17k in pink) and two indices based on GTDB_r86 (GTDB_r86 in orange and GTDB_r86_46k in purple).

We next show that the number of reference genomes rather than strict monophyly of the index database led to increased classification rate. To do so, we compared two indices built using the same reference genomes with (GTDB_r86_8.6k) and without (NCBI_r86) strict monophyletic definitions. When considering all three datasets together, the proportion of unclassified and classified reads/sample were nearly identical using GTDB_r86_8.6k over NCBI_r86 (median increase of +71 classified reads per sample, range of difference from 0 to +2,213 reads/sample, equivalent to a median increase of less than +0.0005% of total reads/sample) (**Figure 1, Table S2, Table S3**), indicating that strict monophyly alone does not substantially affect classification rate. On the other hand, the comparison of classification performance using GTDB_r86 vs GTDB_r86_46k captures the effect of adding reference genomes (28,560 vs 46,006 total genomes, respectively) to two similarly monophyletic indices. When compared to GTDB_r86, GTDB_r86_46k produced more classified reads in almost every (402/404) human and environmental sample tested (**Figure 1, Figure S2, Table S2, Table S3**), with median percentage change in classified reads/sample of +2.7% in human samples (range of −5.6% to +18.7%), +3.2% reads/sample in marine metagenomes (range of +1.8% to +5.0%) and +2.1% (range of +1.9% to +2.3%) in soil metagenomes.

In human samples, a median of 8.6% of total reads/sample (range of 0.9% to 31.6%) (**Table S2**) remained unclassified even when using our best corrected index GTDB_r86_46k. To investigate what these remaining unclassified reads are, we reclassified them using a pre-computed index based on the nucleotide (nt) database of NCBI, which excludes any whole genome sequence from the WGS or RefSeq databases but includes sequences from all taxonomic domains of life (results in **Figure S4**). The large majority of these reads remained unclassified (∼8.5% of total reads/sample); a substantial proportion were attributed to eukaryotic (∼0.86% of total reads/sample) and viral (∼0.16% of total reads/sample) taxa, which are not included in the custom indices used in this study (**Figure S4**). Notably, ∼1.1% of all reads/sample were still attributed to Bacteria and Archaea (**Figure S4**). As the nt database of NCBI excludes reference genomes from WGS and RefSeq, this classification reflects either the presence of genomic fragments in the nt database that are not in WGS of RefSeq, or that these reads mapped to rarer genomic variants that were not included in the 46,006 representative genomes from the GTDB_r86_46k index. As a very small fraction of these unclassified reads were prokaryotic, this result suggests that the GTDB_r86_46k index is much more likely to capture and classify most accessible prokaryotic reads from human metagenomes than default methods.

### Classification to lower taxonomic ranks is increased and more accurate using larger indices

The interpretation of metagenomics data often focuses on lower taxonomic levels, typically genus- and species-level. We compared the taxonomic levels of lowest-common-ancestor (LCA) read classification between the different indices (**Figure 1D-F, Table S4-S5**). The observed trend in all three datasets was that as indices included more reference genomes, more reads were classified to genus and species level (**Figure 1D-F, Table S4-S5**). In particular, GTDB_r86_46k index showed a greater proportion of reads from human samples classified to genus (median increase of +387.2% in reads/sample; range of −22.4% to +3914.7%) and species level (median increase of +44.4% reads/sample; range of −31.6% to +371.7%), as compared with the default NCBI_r86 index (**Figure 1D, Table S4-S5**). For marine samples, corresponding median increases using GTDB_r86_46k were +503.4% reads classified to genus/sample (range of +68.1% to +1124.9%) and +269.2% reads classified to species/sample (range of +64.3% to +567.2%) (**Figure 1E, Table S4-S5**); and +113.4% reads classified to genus/sample (range of +98.8% to +140.0%) and +90.9% reads classified to species/sample (range of +71.4% to +114.8%), for soil samples (**Figure 1F, Table S4-S5**).

Interestingly, of the two best performing indices, GTDB_r86 (built with almost 18,000 less reference genomes than GTDB_r86_46k) classified a median of −28.1% less reads/sample (range of −70.3% to +70.7%) at the genus level, but a median of +3.7% more reads/sample (range of −17.1% to +28.7%) more reads at the species level than GTDB_r86_46k in human samples (**Figure 1D-F, Table S4-S5**). A similar trend was observed in marine and soil samples (**Table S4-S5**). This is likely because the larger the index, the greater the likelihood it includes genomes from two different species that share genes via recent horizontal transfer, which renders those gene sequences ambiguous at the species level so that they can be attributed to their LCA only. In this way, the largest index GTDB_r86_46k can be considered to offer a more accurate representation of taxonomic classification, with ambiguous reads being accurately attributed to the LCA rather than erroneously to a single species.

### The specific composition of corrected indices affects classification performance and detection levels of specific taxa

Unsurprisingly, the specific composition of reference genomes in custom indices affected classification performance. To demonstrate this, we expanded the default NCBI_r88 index by increasing the coverage of 70 human-associated bacterial genera (including pathogens) by 6,819 reference genomes and also correcting monophyly for these genera (to produce the NCBI_r88_Human17k index; **Table 1, File S1**). This expansion of the index from NCBI_r88 had a significantly greater impact on the overall read classification rate for the human metagenomes (mean increase of +44.3% reads/sample; mean range of +0.5% to +249.3%) compared to the environmental metagenomes (mean of +21.2% and +7.0% reads/sample for marine and soil samples, respectively; *p*<0.0001, D=0.3355; Kolmogorov-Smirnov test on human vs. environmental per-sample increase percentage distributions) (**Table S3-S4**). The effect was also clear at both genus and species levels, with mean increases of +181.1% (genus) and +36.4% (species) in human samples compared to mean increases of +28.8% and +10.0% (genus), and +19.2% and +6.5% (species) in marine and soil samples, respectively (**Figure 1D-F, Table S3-S4**).

Specifically, we also observed that 63/70 (90%) of the genera expanded in the NCBI_r88_Human17k index could be classified and detected in human metagenomes at a higher level using this index compared to the default NCBI_r88 index (**Figure S5**). *Pseudomonas, Enterobacter, Butyrivibrio, Lactobacillus, Alistipes, Moraxella, Parabacteroides* and *Faecalibacterium* were amongst the genera with the most significant improvement in detection levels using the expanded index database in HMP metagenomes (**Figure S5**). The detection levels for common human pathogens, including *Yersinia, Clostridium, Helicobacter* or *Acinetobacter*, were also improved when using NCBI_r88_Human17k (**Figure S5B**). In human samples, up to 20% of all reads that remained unclassified using NCBI_r88 but that could be classified using NCBI_r88_Human17k belonged to *Prevotella* and *Bacteroides*, the rest being attributed to a variety of other genera (**Figure S5C, S5D**). When examining particular species of interest, the detection of *Lactobacillus crispatus* in vaginal samples, *Haemophilus parainfluenzae, Campylobacter concisus* and *Campylobacter showae* in buccal and throat samples, were particularly improved by the use of the corrected NCBI_r88_Human17k index, along with numerous distinct species of *Prevotella, Bacteroides* and *Alistipes* in samples from various body sites (**Figure S5C, S5D**). Our results demonstrate that increasing the number of reference genomes for specific genera of interest can substantially improve their detection levels.

### Impact of metagenome-assembled genomes on classification performance

The recently published GTDB taxonomic system (release 86.0) includes 3,087 metagenome-assembled genomes (MAGs) in the taxonomic redefinition of the prokaryotic tree of life [28]. We assessed whether the addition of these potentially new taxa to a reference index improved metagenomic classification on the human, marine and soil test datasets. The addition of 3,087 MAGs to GTDB increased the proportion of reads classified by mean +0.72% (human), +0.63% (marine) and +0.51% (soil) (GTDB_r86_noMAGs vs GTDB_r86 index; **Figure 1, Figure S2, Table S2-S3**). These results show that adding MAGs to index databases can in principle increase classification performance. However, this increase was limited in our test, likely because the MAGs included in GTDB release 86.0 do not capture many novel sequences from the microbiomes analysed (human, marine, and soil).

### GTDB-based species definitions affect taxonomic composition, abundance and diversity metrics, downstream analyses and interpretation from metagenomic studies

The use of corrected indices had a substantial effect on downstream metagenomic analyses. We compared the 30 most abundant taxa for HMP samples at the family, genus and species levels, from classifications using NCBI_r88, NCBI_r88_Human17k and GTDB_r86_46k (**Figure 2, Figure S6**). A total of 19 (63%) families, 15 (50%) genera and 7 (23%) species appeared in the top 30 taxa using all three indices. Thus, the higher the taxonomic order examined, the more agreement across index databases (**Figure 2A, Figure S6**). Notably, even for taxa in the top 30 using all three indices, the order of abundance varied substantially (**Figure 2B, Figure S6**). Some of this variation was attributable to many taxa having been reclassified and renamed in the larger, monophyly-corrected databases (particularly GTDB_r86_46k, see yellow bars in **Figure 2A**). The increased taxonomic granularity within the GTDB system sometimes led to previously common taxa being divided and redefined as multiple different sub-lineages, each with a distinct taxon name. However, there were also differences in the relative abundances of top 30 taxa that were not explained by this (**Figure S7**). For example, the relative abundance rank of families *Porphyromonadaceae* and *Corynebacteriaceae* were reversed using NCBI_r88 vs. GTDB_r86_46k, as were the genera *Lactobacillus* and *Bifidobacterium*, and the species *Bacteroides fragilis* and *Bacteroides thetaiotaomicron* (**Figure S7**).

**Figure 2.**
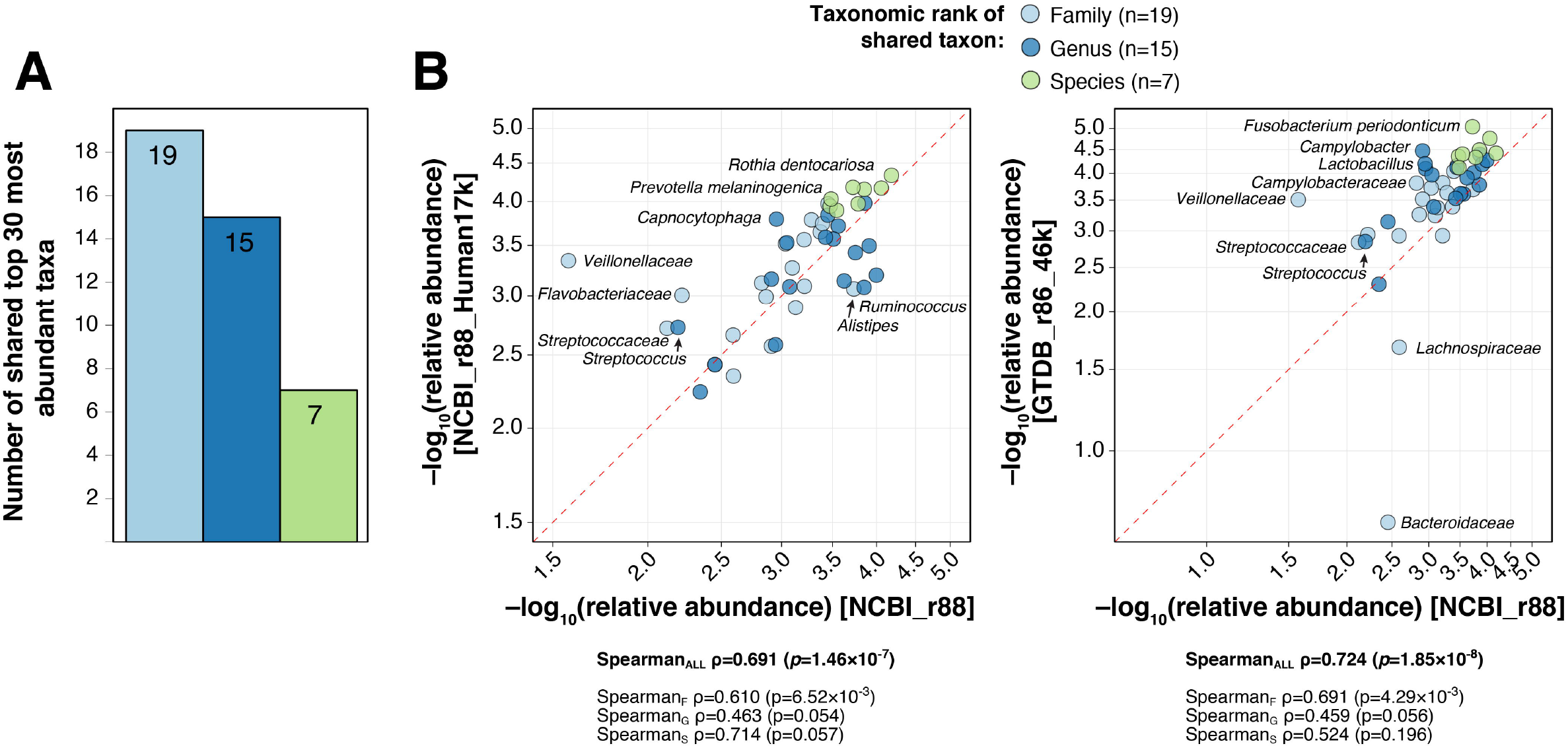
Effect of index database correction on metagenomic composition. (A) Number of shared top 30 most abundant families, genera and species after classification of 98 HMP samples using default NCBI_r88 index and corrected NCBI_r88_Human17k and GTDB_r86_46k indices. (B) Comparisons of relative abundances (−log_10_ scale) between the default NCBI_r88 classification and NCBI_r88_Human17k classifications (left) and GTDB_r86_46k (right) for taxa in the top 30 most abundant of all three classifications (19 families, 15 genera and 7 species). To assess changes in rank order consistency between the classifications, Spearman’s rank correlation coefficient, and the associated p-value are shown for both comparisons of NCBI_r88_Human17k and GTDB_r86_46k classifications with NCBI_r88 for all taxa, and at each taxonomic ranks.

Alpha diversity (within-sample diversity), which has been associated with various phenotypes in different microbiomes [6, 31–33], is estimated directly from taxonomic composition data and therefore showed significant differences between indices. We compared three alpha-diversity metrics at the genus level (observed genus richness, genus evenness and Shannon index at the genus level), calculated from taxonomic composition tables summarised at the genus level based on classifications of the same test data sets but using seven different index databases (**Figure 3**). As expected, the large GTDB-based indices showed a much higher richness, but also had an effect on the evenness of genus distribution, especially in marine metagenomes, which affected Shannon diversity index distribution (**Figure 3**). Notably the effect of index database on alpha diversity values varied between samples, with some increasing in value and others decreasing. In some cases these differences were substantive enough to alter the results of statistical tests for difference in alpha diversity between samples from different body sites (**Figure 3B, Table S6**). For example, in our subset of the HMP dataset, faecal samples were found to have significantly lower Shannon diversity than buccal samples when using the NCBI_r86 index (median of 1.44 [IQR 1.03-2.25] vs median of 2.41 [IQR 2.02-2.70] respectively, p=0.027) (**Table S6**). A similar result was obtained using NCBI_r88 index (**Table S6**). However no such differences were found between faecal and buccal samples when Shannon diversity was calculated using any of the GTDB-based indices (median of 2.09 [IQR 1.62-2.83] vs median of 2.59 [IQR 2.16-2.90] respectively, p=0.999 using GTDB_r86_8.6k) (**Table S6**). The situation was reversed when comparing Shannon diversity for faecal and skin samples, with significant differences obtained using GTDB_r86_8.6k (median of 2.09 [IQR 1.62-2.83] vs median of 0.81 [IQR 0.63-1.12], p=0.001) but not using NCBI_r86 (median of 1.44 [IQR 1.03-2.25] vs median of 0.77 [IQR 0.63-1.12], p=0.965) (**Table S6**).

**Figure 3.**
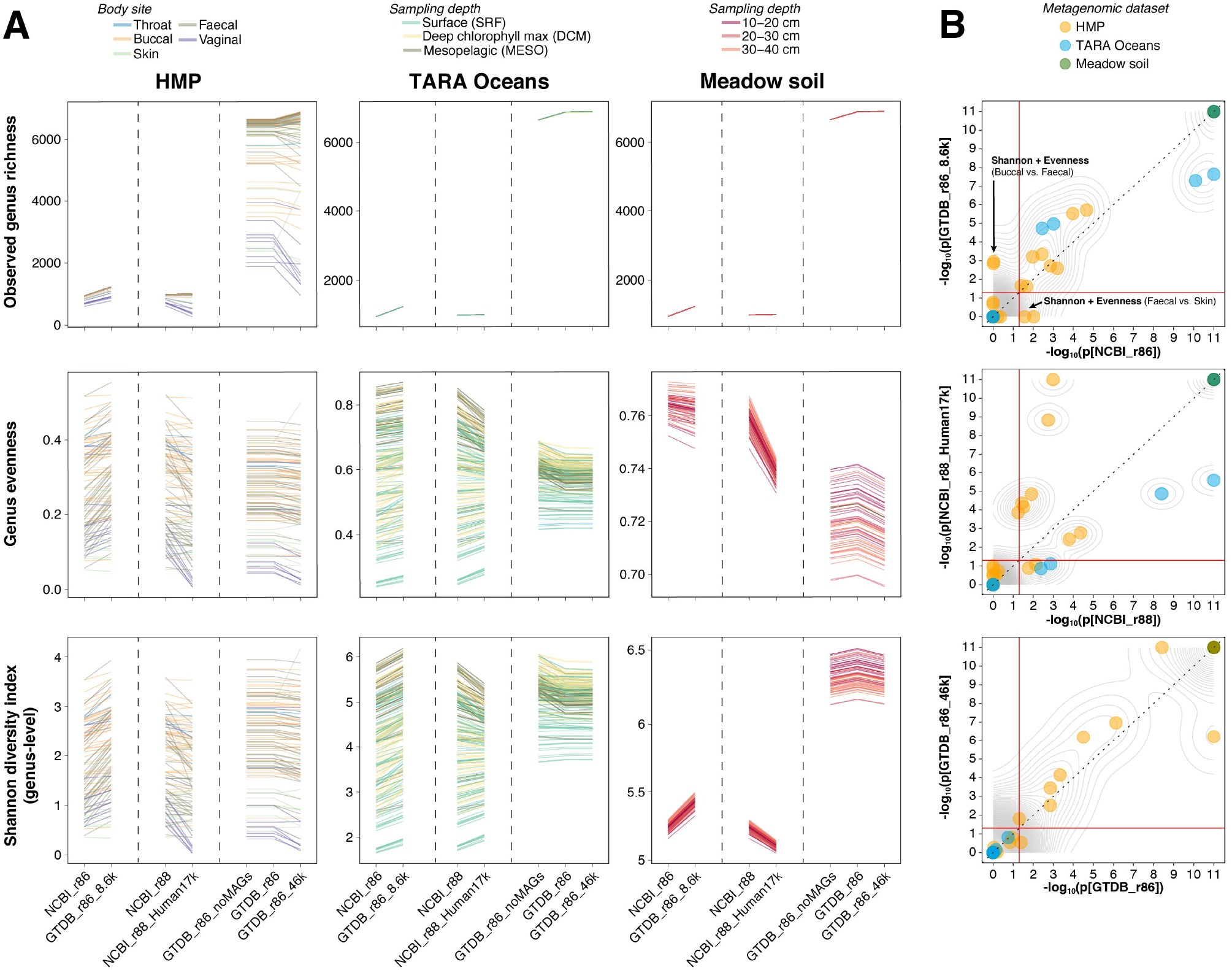
Using corrected indices to classify metagenomes affects measures of alpha diversity. (A) The values of three measures of alpha diversity (observed genus richness, genus evenness and Shannon diversity index at the genus level) for each metagenomic sample from three datasets (HMP subset, TARA Oceans, Meadow soil samples) are shown. Three specific comparisons of values are presented, between NCBI_r86 and GTDB_r86_8.6k, between NCBI_r88 and NCBI_r88_Human17k and between GTDB_r86, GTDB_r86_noMAGs and GTDB_r86_46k. Each sample is represented by a line coloured by isolation phenotype. Statistical comparisons of distributions presented in this panel are shown in Table S6. (B) Effect of classification index on alpha diversity metrics comparisons between groups. The scatter plots compare the significance of ANOVA tests on all alpha-diversity measures for each of three comparisons, P-values using Dunn’s multiple testing (with Holm’s correction). The dotted lines represent proportionality for which p-values are identical for classifications using both indices, the red lines denote the p-value threshold of 0.05 (−log_10_=1.301) for each index.

We also examined the effect of index database choice on beta-diversity, or between-sample diversity assessed by calculating Bray-Curtis dissimilarity between groups of samples from different sources (**Figure S8, S9, S10, Table S7**). The effect on beta-diversity was more subtle than for alpha diversity, with the large GTDB indices yielding greater distance estimates between groups of samples that were already dissimilar using default methods (dissimilarity above 80%; **Figure S8, S9**), but did not significantly alter the overall clustering patterns (**Figure S10**).

## Discussion

Considerable efforts have been made to improve methods for detection of taxonomic and functional markers in complex metagenomic samples, including increasing sequencing depth, optimising classification algorithms and developing more accurate *de novo* metagenome assembly tools. In this study, we showed that the index database is a major source of variation in classification performance and has significant ramifications for downstream analyses, which may be substantive enough to change study conclusions (e.g. alpha diversity). Commonly utilised index databases lead to sub-optimal taxonomic classification, with a minority of some read sets being classified. Increasing the number of phylogenetically consistent reference genomes in an index database in either a broad or targeted manner had consistently positive effects on increasing the proportion of reads classified (sometimes several fold higher) and classification to greater taxonomic resolution. To facilitate metagenomic analyses without the need for deeper sequencing or *de novo* assembly, we make freely available these improved index databases (https://github.com/rrwick/Metagenomics-Index-Correction) for two commonly-used classifiers, Centrifuge and Kraken2 and the tools to construct them as NCBI RefSeq and GTDB expand.

We found that large indices built using recently developed and largely phylogenetically-coherent taxonomic species definitions, such as GTDB [28], greatly increased the number of classified reads. Our results suggest that more coherent taxonomic definitions and accurate taxonomic boundaries, such as those proposed within GTDB, may improve statistical power and biological interpretation of subsequent results, particularly those for compositional and diversity analyses (summarised in **Figure 4**). This results in greater taxon granularity, i.e. smaller, more discrete clades of similar phylogenetic depth than commonly known phylogroups, which increases classification accuracy and may improve downstream applications, such as association analysis for particular traits. For example, in microbiome-wide association studies using large cohorts, a weak association with a poorly-defined lineage may be caused by a strong association with a well-defined subset of the poorly-defined lineage (**Figure 4**). Furthermore, at a fixed confidence level, increasing the classification rate of a metagenomic sample offers a more accurate representation of its microbial diversity and may, as we have shown, affect study conclusions. As such, the approach we propose here facilitates improved metagenomic analysis across the full spectrum of sequencing depths. In particular, our results may facilitate “shallow sequencing” metagenomics [34] by maximising the extraction of taxonomic information from samples sequenced at lower depth, thus enabling more cost-effective comparison of thousands or tens of thousands of samples in large-scale metagenomic and multi-omics studies. Lastly, our study shows the importance of consistency in index database when comparing results across studies. Differences in reference genomes and taxonomic coherence may introduce artefacts when integrating metagenomic data across studies, and therefore care should be taken when performing combined or meta-analyses.

**Figure 4.**
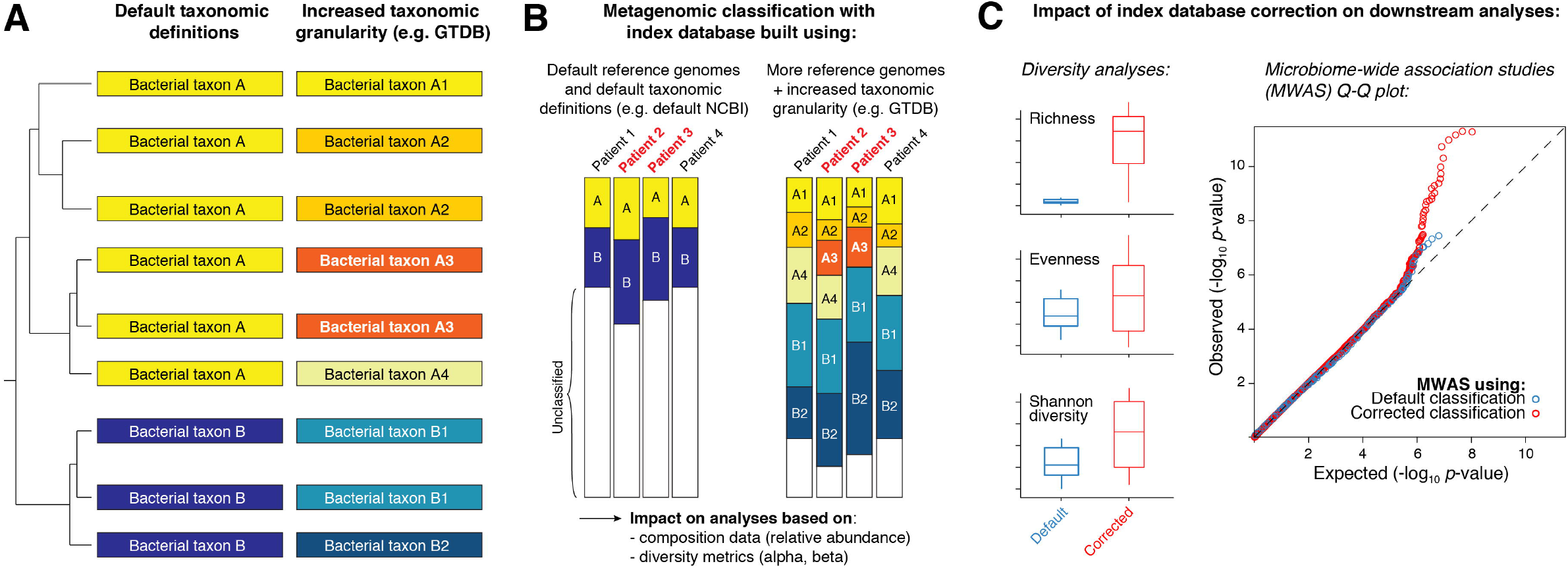
Increased taxonomic granularity in classification indices can improve the interpretation of microbiome-wide association studies. (A) Increased taxonomic granularity is defined here by the accurate redefinition, splitting and merging of phylogenetically-coherent strictly-monophyletic lineages, as performed using GTDB. In this example, taxon A is split into taxa A1, A2, A3 and A4, and taxon B is split into B1 and B2. (B) Example classification using two index databases, a smaller number of reference genomes with polyphyletic definitions (left) and a larger number of reference genomes with monophyletic definitions (right). (C) Example effects of index database correction on downstream analysis involving alpha-diversity metrics (left) or for microbiome-wide association studies (right).

## Material and Methods

### Description of corrected index databases

Seven different indices, ranging in size from 8674 to 46,006 complete genomes, were built to compare the effect of various factors on metagenomic classification performance (**Table 1**). As the focus of this study was not to compare the performance of specific metagenomic classifier tools themselves, but rather to evaluate the impact of custom indices on metagenomic classification, we picked a recently developed classifier, Centrifuge [19], on the basis of an easy-to-use index customisation pipeline, fast metagenomic classification performance and lower RAM usage than alternative tools. Centrifuge allows for the building of custom indices (via the *centrifuge-build* indexer), taking as input a set of sequences with taxonomic labels and a ranked taxonomic tree describing the relationships between those labels.

The NCBI_r86 and NCBI_r88 indices were built from the default collections of complete bacterial and archaeal genomes from NCBI RefSeq releases 86 (n=8,674 genomes) and 88 (n=10,089 genomes), respectively, using the NCBI taxonomy tree. The NCBI_r88_Human17k index was based on the NCBI_r88 index, with the addition of 6819 further reference genomes from NCBI GenBank plus manual curation of the taxonomy for 70 common human commensal and pathogenic bacterial genera (**File S1**). We used the Bacsort pipeline (https://github.com/rrwick/Bacsort) to manually curate the taxonomy within each of these 70 genera to enforce strict monophyly. We also built four indices using the GTDB taxonomic system (**Table 1**). GTDB is based on curation of >125,000 whole genome sequences sourced from NCBI RefSeq and metagenome-assembled genomes (MAGs); but with taxonomic labels and tree re-defined based on phylogenetic relationships inferred from the concatenation of 120 proteins and enforcing strict monophyly [28]. We built the GTDB_r86 index from the default GTDB release 86 set of 28,941 dereplicated bacterial and archaeal genomes representative of the GTDB taxonomy [28], “dereplication” being defined as the selection of reference genomes representative of phylogenetic similarity clusters [28]. In the original GTDB publication and website, two genomes were found to be “replicates” when a set of conditions were met, typically when their Mash distance was ≤0.05 (∼ANI of 95%) [4]. The GTDB_r86_8.6k index was built using the exact same 8,674 complete reference genomes as for NCBI_r86, but using the taxonomic labels and trees assigned by GTDB. The GTDB_r86_noMAGs index was built exactly like GTDB_r86, but excluding all 3,087 metagenome-assembled genomes (MAGs) identified in GTDB release 86. Finally, the GTDB_r86_46k index was built using a lower Mash threshold for dereplication than in the default GTDB_r86 set. Specifically, this index included a total of 46,006 reference genomes (18,634 more than GTDB_r86), each representative of similarity clusters defined using a Mash [10] distance threshold of ≤0.005 (∼ANI of 99.5%). The tax_from_gtdb.py and dereplicate_assembly.py scripts are available in https://github.com/rrwick/Metagenomics-Index-Correction with instructions.

### Metagenomic datasets

We used a total of 404 publicly available metagenomes representing a variety of commonly-studied environments: human body sites, marine and soil environments (**Table S1, Figure S1**). Human samples were from the WGS-PP1 study of the Human Microbiome Project (HMP) and obtained through the HMPDACC.org website [9]. HMP samples were chosen with the following considerations: we kept a representative proportion of each body site represented in the WGS-PP1 study, we did not subset a body site source with less than five samples, and we excluded samples with a high (>90%) proportion of low quality reads and samples with low sequencing depth. A total of 98 representative samples were selected, corresponding to ∼9.2% (n=98/1067) of the HMP WGS-PP1 study. A total of 246 marine metagenomic samples were isolated from a range of locations in epipelagic and mesopelagic waters around the world as part of the TARA Oceans survey [35, 36], and were downloaded from the EBI repository (study MGYS00002008; BioProject PRJEB1787). A total of 60 soil metagenomes were sampled in a recent study from meadows ground at various depths [37], and were obtained from NCBI BioProject PRJNA449266. Accessions for all readsets are listed in **Table S1**.

### Assessment of metagenomic classification performance

For all classifications, we ran Centrifuge version 1.0.4 [19] on a Linux x86 cluster with 16 cores and 128 GB of RAM allocated for each sample classification. The run time ranged from 11 to 45 minutes per metagenomic sample, depending on the index used for classification and the sequencing depth of the sample. Classification reports were built from the resulting output files using the *centrifuge-kreport* tool, and reports were visualised and exported using Pavian version 0.8.1 [38] and custom-made scripts, available at https://github.com/rrwick/Metagenomics-Index-Correction.

Classification performance was assessed by first comparing the number of unclassified and classified reads per sample for each index database used. This provides an unambiguous way to measure how much of the total microbial information present in each sample can be classified. We also compared the taxonomic ranks to which reads were assigned using each index. It should be noted that the NCBI prokaryotic taxonomic system includes many additional and ambiguous taxonomic ranks that are not present in GTDB, such as “subphylum”, “infraclass”, “superclass”, “subtribe” or “strain”. To make results comparable between taxonomic systems, reads were always attributed and reported to the LCA of the standard ranks: phylum, class, order, family, genus and species.

A pre-compiled index based on the nucleotide (nt) database is available from the Centrifuge website (http://www.ccb.jhu.edu/software/centrifuge/, compiled on 03/03/2018). This database includes all traditional divisions of GenBank, EMBL and DDBJ, and thus includes eukaryotic and viral sequences in addition to prokaryotes. However, the nt database excludes the WGS section of GenBank, which should have a negative impact on the determination of accurate species-specific microbial markers. Accordingly, we observed that the classification of 10 random HMP metagenomes using nt resulted in more unclassified reads than when using GTDB_r86_46k (data not shown). To investigate the origin of the reads which were unclassified by the GTDB_r86_46k index (the best-performing custom index in this study), we reclassified them using the nt database.

Finally, we assessed the effect of using different indices on commonly-used ecological diversity metrics. The calculation of alpha and beta diversity estimates (observed genus richness, genus evenness, Shannon diversity and Bray-Curtis dissimilarity at the genus level) was performed using the R package *phyloseq* version 1.24.2 [39].

### Custom scripts and pre-computed index databases availability

A collection of scripts used to prepare, compare and analyse Centrifuge classifications using custom index databases, either based on default NCBI or GTDB taxonomic systems, is available at: https://github.com/rrwick/Metagenomics-Index-Correction with instructions. Pre-computed versions of the NCBI_r88_Human17k and GTDB_r86_46k indices suitable for use with Centrifuge [19], Kraken1 [18], Kraken2 (https://ccb.jhu.edu/software/kraken2/) and their variants (KrakenUniq [17], LiveKraken [40]), are freely available to from: https://monash.figshare.com/projects/Metagenomics_Index_Correction/65534.

## Authors’ contributions

GM designed the study, generated performed analyses, interpreted results and was the major contributor in writing the manuscript. RRW helped generating scripts and databases, interpreted results and participated in the writing of the manuscript. SCW helped to generate code and databases. KEH and MI were key contributors to the study design, interpretation of results and the writing of the manuscript. All authors read and approved the final manuscript.

---

**Figure S1.**
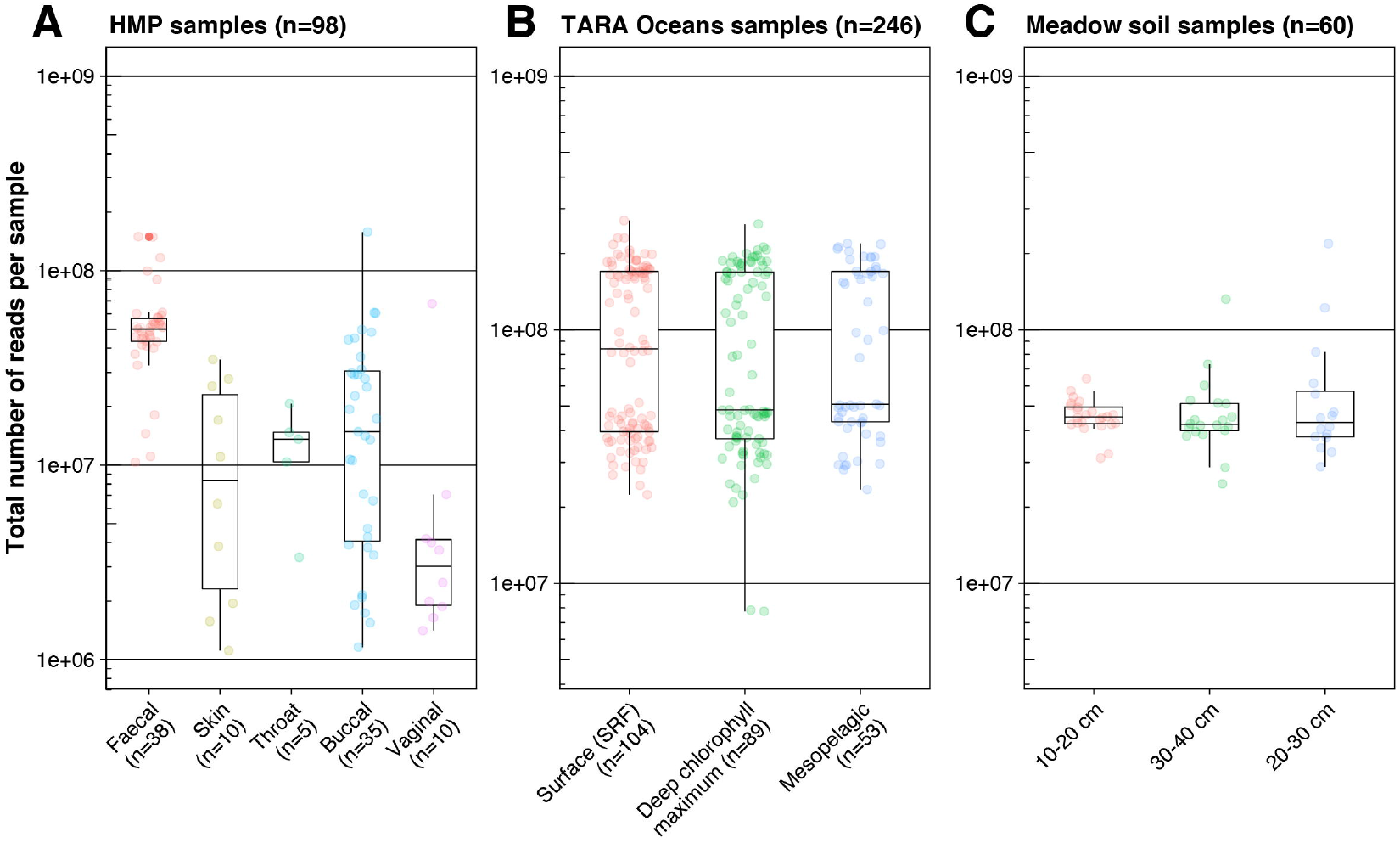
Description of 404 metagenomic samples used in this study. The distribution of the number of reads/sample is shown for 98 human (A), 246 marine (B) and 60 soil (C) samples, according to various sampling information (body site for human samples, and sampling depth for marine and soil samples).

**Figure S2.**
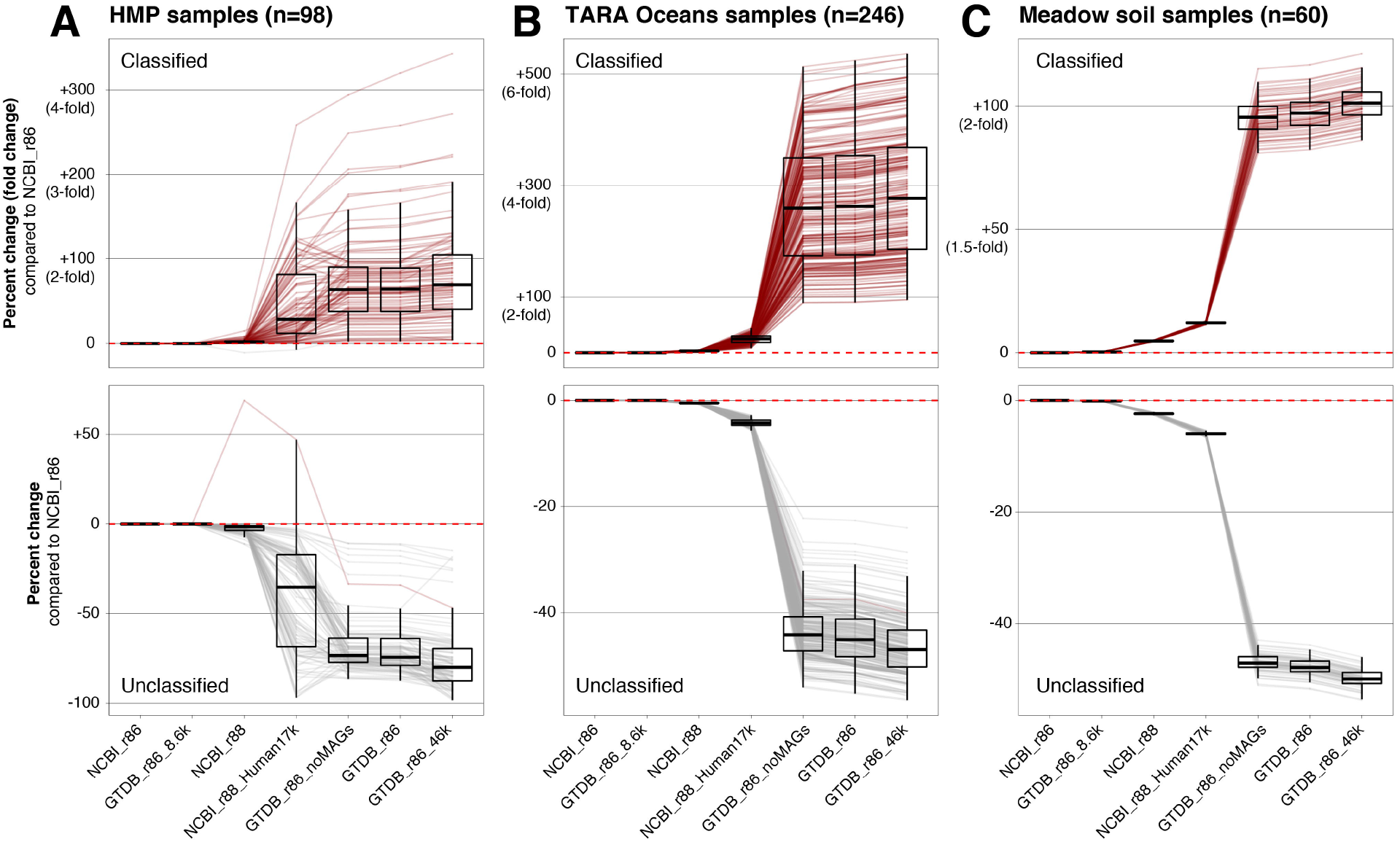
Per-sample change (% and fold-change) in unclassified and classified reads/sample using seven default and corrected NCBI- and GTDB-based indices. Per-sample percent and fold-changes are shown for the three metagenomic datasets: (A) human samples (n=98), (B) TARA Oceans samples (n=246) and (C) meadow soil samples (n=60). Values are normalised to the number of reads unclassified and classified using the default NCBI_r86 index.

**Figure S3.**
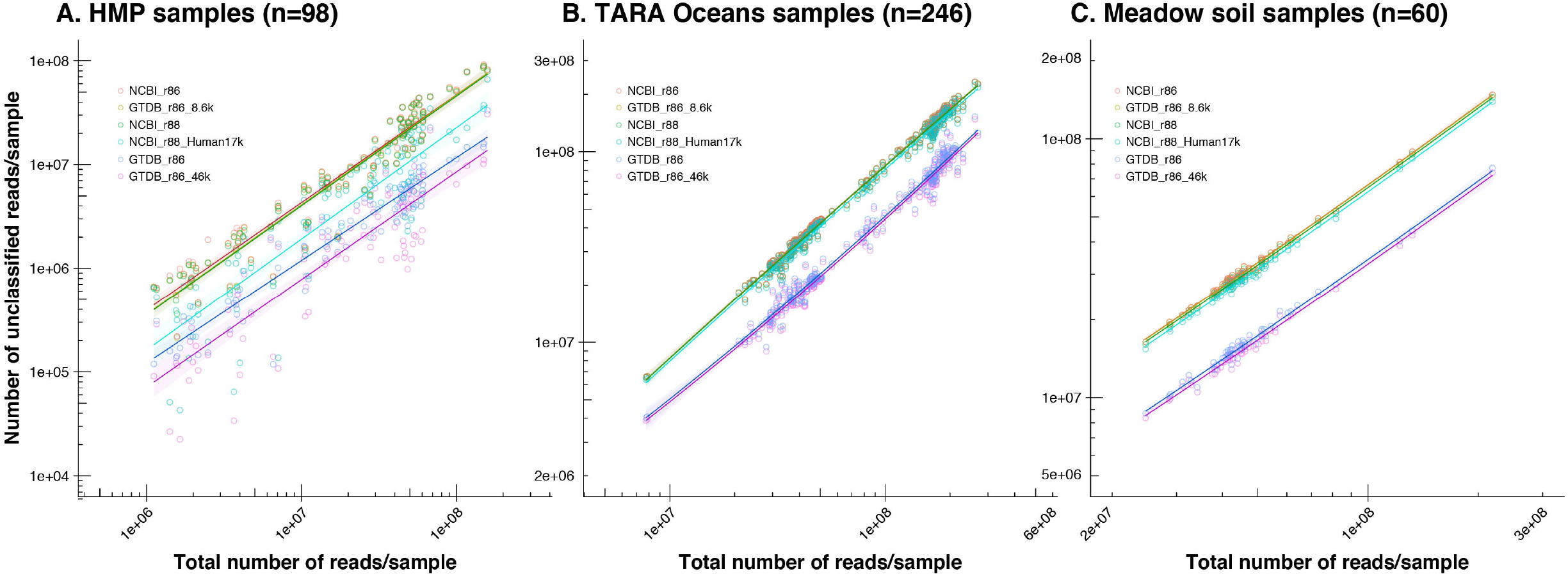
Classification improvement using corrected indices is unaffected by variations in sequencing depth. The total number of reads/sample (a proxy for sequencing depth) was plotted against the number of unclassified reads/sample using 6 default and corrected indices, for human (A), marine (B) and soil (C) metagenomes. The regression line was calculated using a linear model fit (“lm” in ggplot2 *geom_smooth* function) for each index.

**Figure S4.**
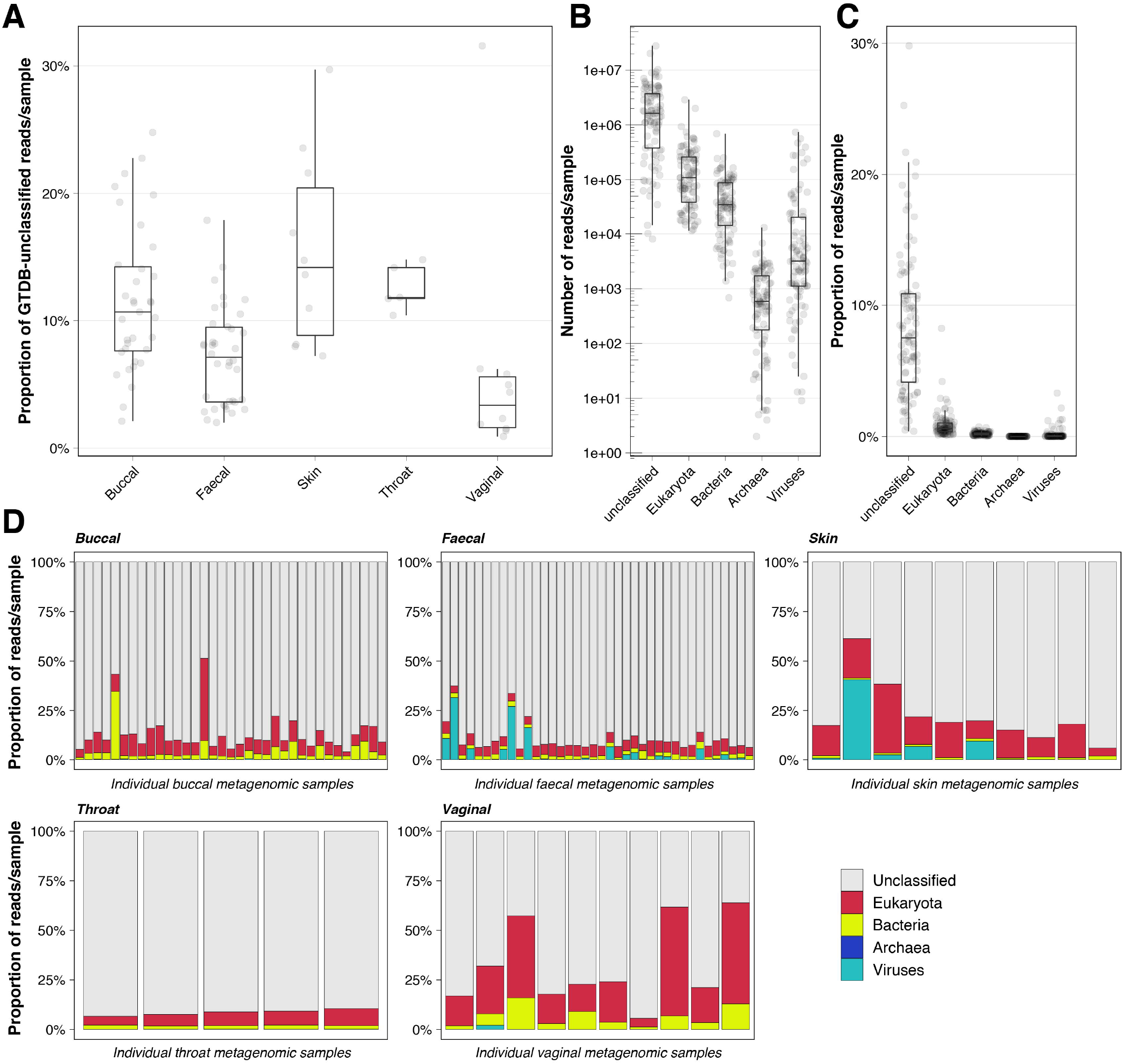
Reads from human metagenomes that remained unclassified using the GTDB_r86_46k index are mostly unknown and eukaryotic. (A) Proportion of reads/sample from HMP samples (n=98) that are unclassified using GTDB_r86_46k, according to body site of isolation; (B) and (C) outcome of re-classification of these specific reads using an index based on the NCBI nucleotide database (nt; pre-computed on the 03/03/2018 and available on the Centrifuge website [http://www.ccb.jhu.edu/software/centrifuge/]) in number of reads/sample (B) and in proportion (C); (D) per-sample breakdown of domain re-classification, showing the proportion of reads attributed to Eukaryota, Bacteria, Archaea and Viruses or unclassified.

**Figure S5.**
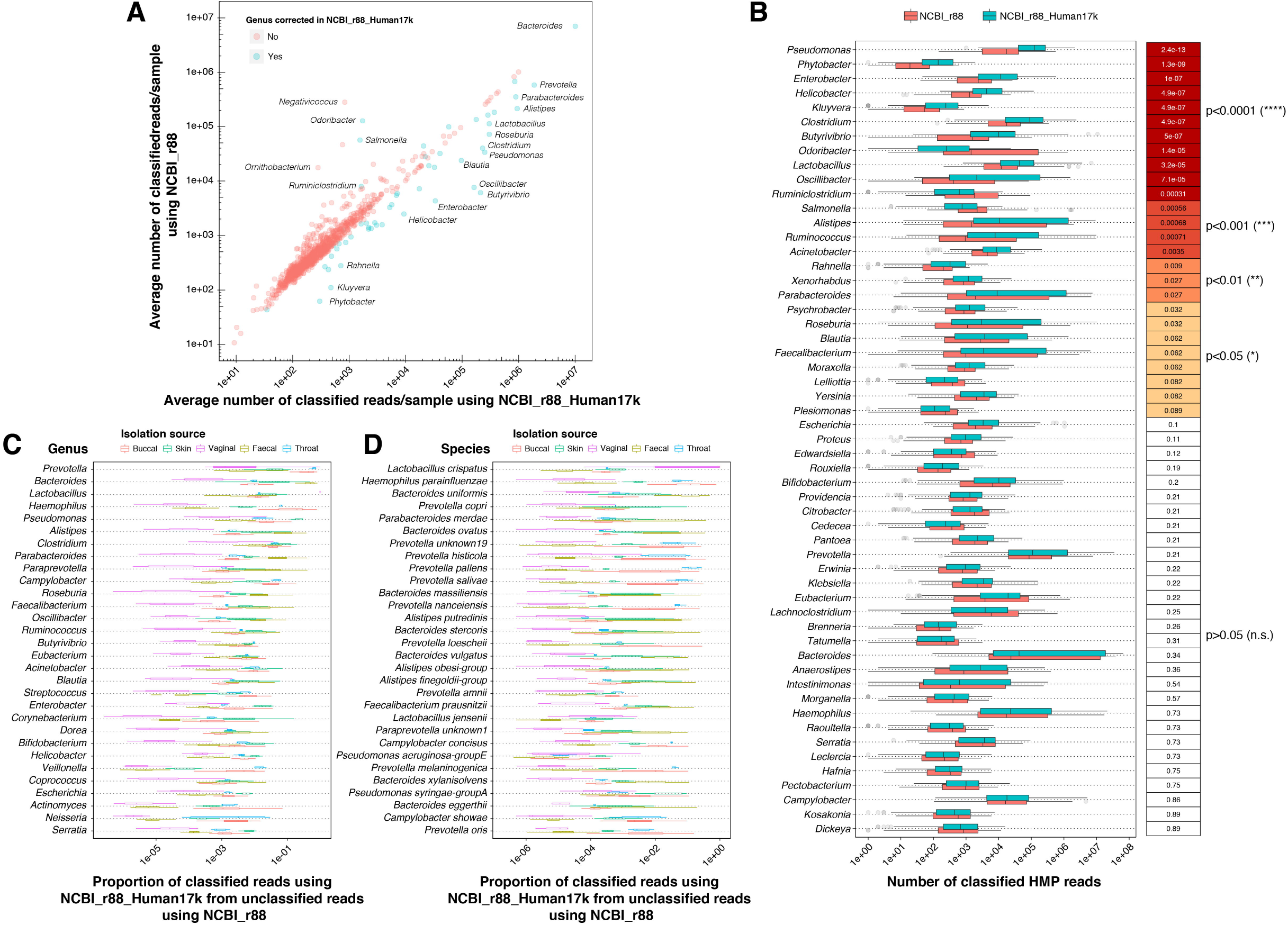
Targeted correction for 70 specific bacterial genera increases their detection levels in human metagenomes. (A) The average number of reads classified to all 1001 different bacterial genera to which at least one read was attributed using the NCBI_r88 and the NCBI_r88_Human17k indices were compared, with the 70 specifically corrected genera were highlighted in blue while the non-corrected genera are shown in red. Any point above the line denotes genera to which more reads were classified using the NCBI_r88 index, while any point below the line denotes genera to which more reads were classified using the NCBI_r88_Human17k index. (B) Number of classified reads/sample using a default index and a targeted correction for 70 specific bacterial genera. The 70 corrected genera are shown, along with their corresponding distribution of the number of classified reads/sample using NCBI_r88 (red) or NCBI_r88_Human17k (blue). The column on the right indicates the p-value and significance thresholds after Wilcoxon signed-rank tests comparing the two indices. Genera with the highest significance in difference are shown in red, orange and yellow, and non-significant differences are shown in white. (C and D) From all reads that were unclassified using NCBI_r88 but classified using NCBI_r88_Human17k, the top 30 genera (C) and species (D) to which these reads were attributed, in proportion, are shown. Boxplots of different colours denote different isolation sources, showing how body sites are differently affected by the index correction.

**Figure S6.**
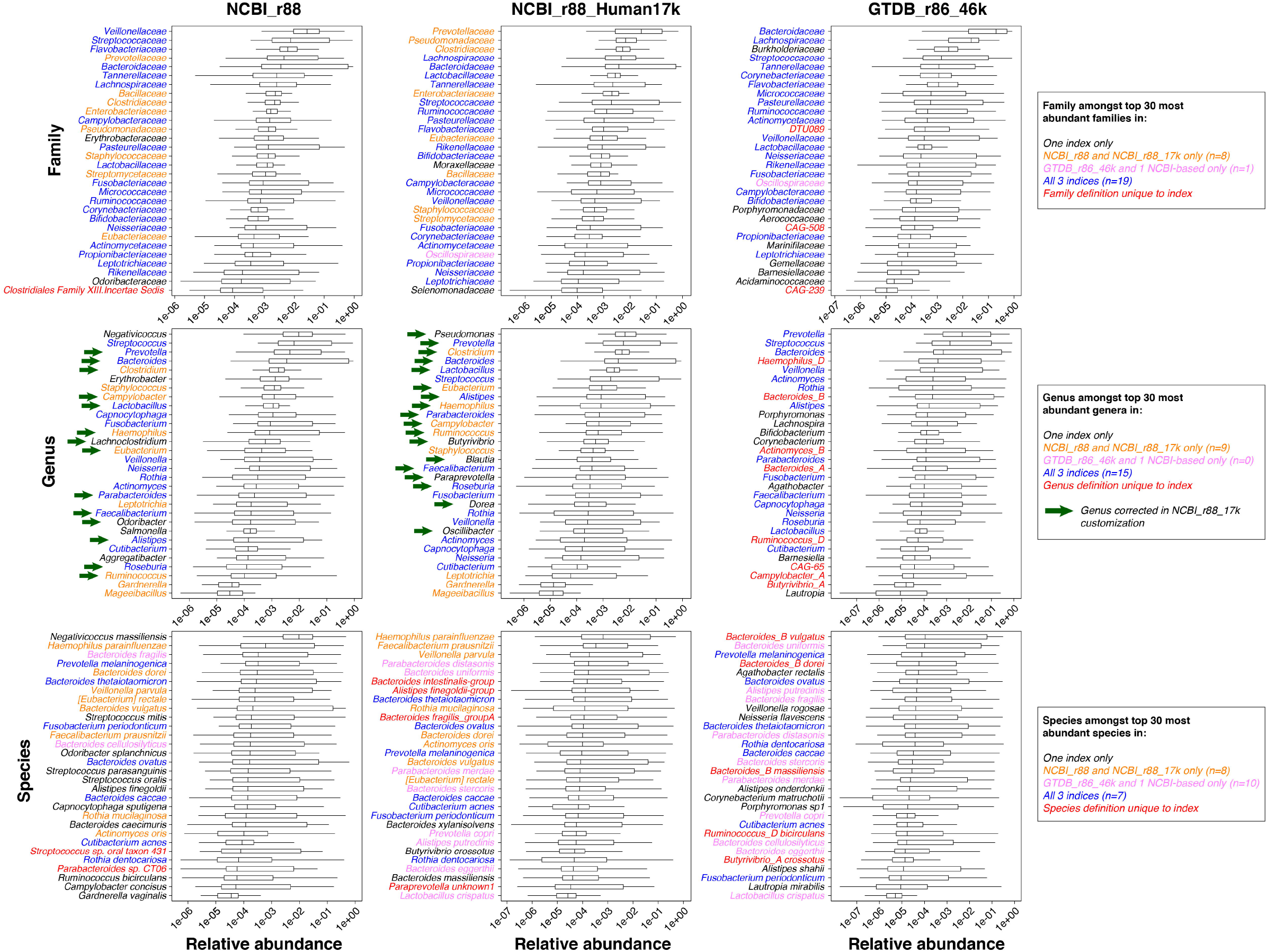
Effect of index database correction on metagenomic compositional data and most abundant taxa. The 30 most relative abundant families, genera and species to which reads were attributed using three indices (default NCBI_r88 and corrected indices NCBI_r88_Human17k and GTDB_r86_46k) are shown in boxplots. The colour of the taxon name in the y-axes denotes whether the same taxon label was found to be present in the top 30 most abundant taxa after classification by all three indices (blue), by NCBI_r88 and NCBI_r88_Human17k and not GTDB_r86_46k (orange), by GTDB_r86_46k and either NCBI_r88 or NCBI_r88_Human17k (pink) or only in one index (black). In red are indicated the taxon definitions that are existing only in one index. For the comparison at the genus level, green arrows indicate whether the corresponding genus has been specifically corrected in the NCBI_r88_Human17k index.

**Figure S7.**
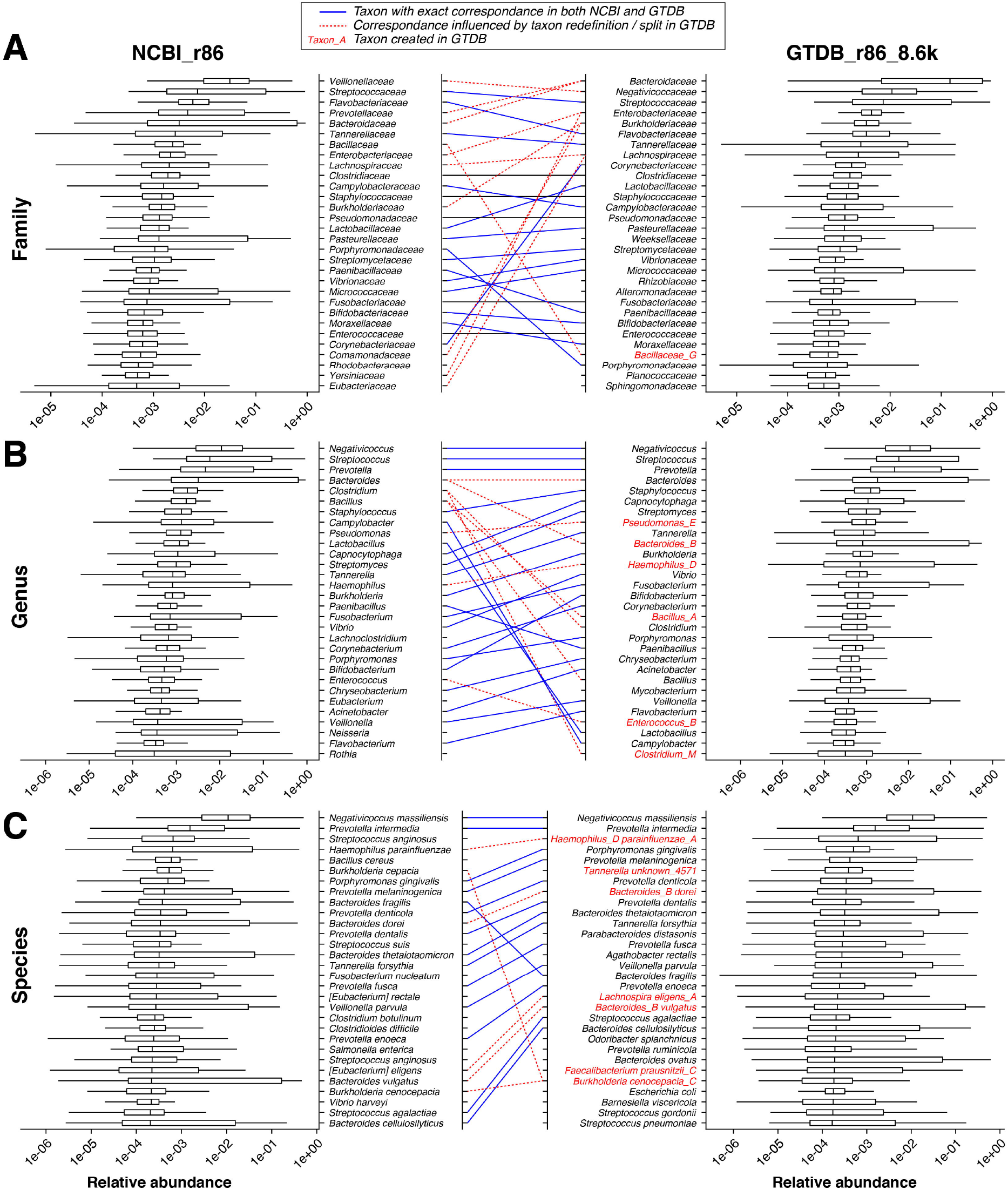
Comparison of metagenomic compositional data and most abundant taxa after classification with indices built from the same reference genomes taxonomically defined using NCBI- or GTDB-based definitions. The 30 most relative abundant families (A), genera (B) and species (C) to which reads were attributed using the NCBI_r86 and GTDB_r86_8.6k indices, built with the exact same set of complete reference genomes from NCBI RefSeq release 86, are shown in boxplots. The correspondence between the top 30 most abundant taxa from the two classifications is reflected by lines between the two plots. The colouring of the lines denote taxa with an exact correspondence in both indices (plain blue) or whether the GTDB redefinition of taxa affected the correspondence (dotted red). The taxa written in red were created in GTDB.

**Figure S8.**
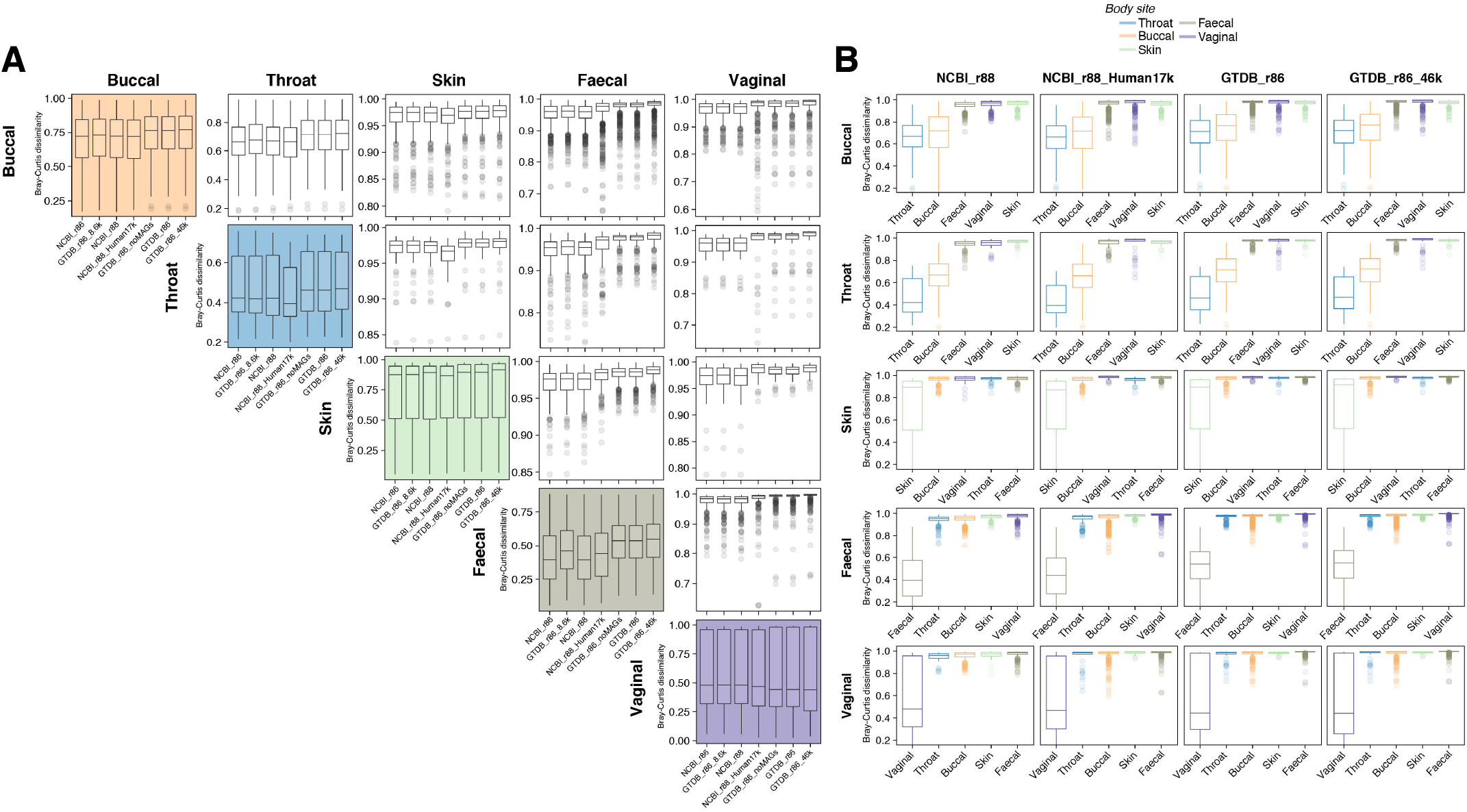
Effect of using corrected indices to classify metagenomes on Bray-Curtis dissimilarity between groups of HMP samples (grouped by body site isolation). (A) Bray-Curtis dissimilarity distributions are shown for pairwise group comparisons between buccal, throat, skin, faecal and vaginal samples of the HMP dataset subset (n=98) used in this study, using seven different classification indices. Coloured panels denote within-group comparisons, white panels denote between-group comparisons. (B) Visualisation of the same data, but ordered to contrast the effect of index database on pairwise group comparisons of Bray-Curtis dissimilarity. Colours represent body sites, similarly to panel A.

**Figure S9.**
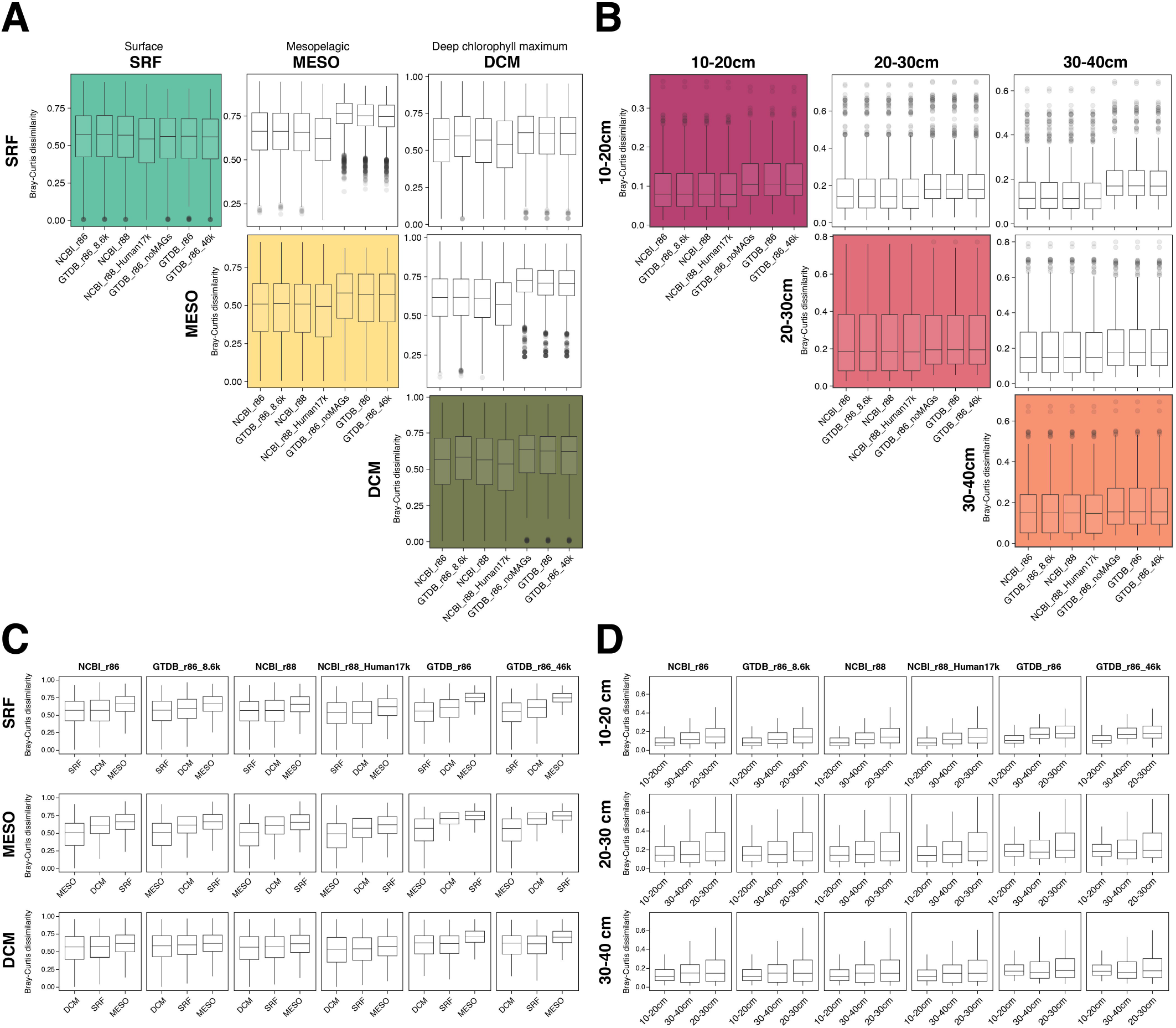
Effect of using corrected indices to classify metagenomes on Bray-Curtis dissimilarity between groups of TARA Oceans and meadow soil samples (grouped by body site isolation). (A-B) Bray-Curtis dissimilarity distributions are shown for pairwise group comparisons between buccal, throat, skin, faecal and vaginal samples of the TARA Oceans dataset (n=246, panel A) and meadow soil dataset (n=60, panel B) used in this study, using seven different classification indices. Coloured panels denote within-group comparisons, white panels denote between-group comparisons. (C-D) Visualisation of the same data, but ordered to contrast the effect of index database on pairwise group comparisons of Bray-Curtis dissimilarity for TARA Oceans samples (panel C) and meadow soil samples (panel D). Statistical comparisons of distributions presented in this figure are shown in Table S7.

**Figure S10.**
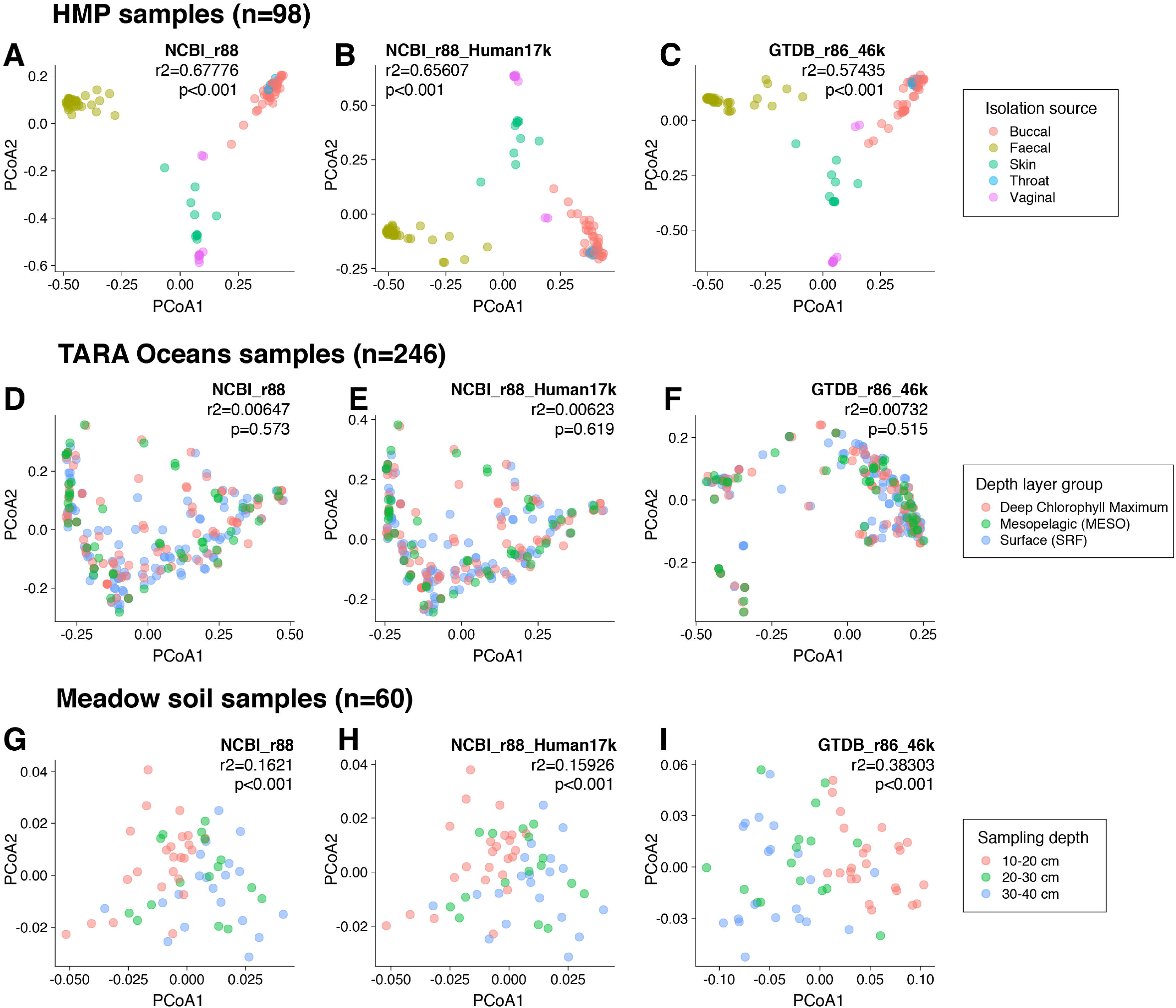
Effect of using corrected indices to classify metagenomes on Bray-Curtis dissimilarity (ordination plots). Between-sample diversity was compared by calculating and ordinating Bray-Curtis distance measures for samples classified using NCBI_r88 (C, H, M), NCBI_r88_Human17k (D, I, N) and GTDB_r86_46k (E, J, O). To compare the effect of indices on beta-diversity, we performed permutational multivariate analysis of variance (PERMANOVA) on the Bray-Curtis distances to measure the association between sample information (such as isolation source for the HMP samples, or sampling depth for the marine and soil samples) and variance within the dataset, indicated in bold in panels C-E, H-J, M-O.

**Table S1. Sample description and accession number for 404 public human, marine and soil metagenomes used in this study.**

**Table S2. Summary of classification outcome for three datasets using seven different index databases.** The median, average, minimum and maximum values of the number and proportion of classified and unclassified reads/sample is shown for every condition. Classifications were performed using Centrifuge version 1.0.4. Detailed per-sample values are shown in Table S3.

**Table S3. Detailed per-sample classification outcome using samples from three datasets, after classification using seven different index databases.** The number and proportion of classified and unclassified reads/sample is shown for every condition. Classifications were performed using Centrifuge version 1.0.4. A summary of these results is shown in Table S2.

**Table S4. Summary of classification outcome at the genus and species level for samples from three datasets using five different index databases.** The median, average, minimum and maximum values of the number and proportion of classified reads/sample is shown for every condition. Classifications were performed using Centrifuge version 1.0.4. Detailed per-sample values are shown in Table S5.

**Table S5. Detailed per-sample classification outcome to genus and species levels using samples from three datasets, after classification using five different index databases.** The number and proportion of classified reads/sample is shown for every condition. Classifications were performed using Centrifuge version 1.0.4. A summary of these results is shown in Table S4.

**Table S6. Influence of index correction on the significance of differences between alpha diversity of isolation phenotype groups.** Comparison of significance (p-values) from Dunn’s multiple testings with Holm correction after ANOVA on three alpha diversity metrics comparisons (observed richness, evenness and Shannon diversity index) between isolation phenotype groups in three metagenomic datasets (body site for HMP samples, depth of sampling for TARA Oceans and meadow soil samples) after classification with six index databases. Specifically, the alpha diversity metric distribution between isolation group pairs were compared after classifications with NCBI_r86 and GTDB_r86_8.6k, NCBI_r88 and NCBI_r88_Human17k and GTDB_r86, GTDB_r86_noMAGs and GTDB_r86_46k.

**Table S7. Influence of index correction on the significance of differences between beta diversity (Bray-Curtis dissimilarity) calculated between isolation phenotype groups.** Comparison of significance (p-values) from Dunn’s multiple testings with Holm correction after ANOVA on Bray Curtis dissimilarity comparisons between isolation phenotype groups in three metagenomic datasets (body site for HMP samples, depth of sampling for TARA Oceans and meadow soil samples) after classification with four index databases. Specifically, comparisons were between NCBI_r88 and NCBI_r88_Human17k and GTDB_r86 and GTDB_r86_46k.

**File S1. Description of the NCBI_r88_Human17k index database creation.** Pre- and post-correction Newick and XML phylogenetic trees built on hybrid FastANI/Mash distances for 9928 genomes from 70 genera of interest (*Acinetobacter, Alistipes, Anaerostipes, Atlantibacter, Bacteroides, Barnesiella, Bifidobacterium, Blautia, Brenneria, Buttiauxella, Butyrivibrio, Campylobacter, Cedecea, Citrobacter, Clostridium, Coprococcus, Dickeya, Dorea, Edwardsiella, Enterobacter, Erwinia, Escherichia, Eubacterium, Faecalibacterium, Haemophilus, Hafnia, Helicobacter, Intestinimonas, Izhakiella, Klebsiella, Kluyvera, Kosakonia, Lachnoclostridium, Lactobacillus, Leclercia, Lelliottia, Mangrovibacter, Moraxella, Morganella, Nissabacter, Odoribacter, Oscillibacter, Pantoea, Parabacteroides, Paraprevotella, Pectobacterium, Phascolarctobacterium, Phytobacter, Plesiomonas, Prevotella, Proteus, Providencia, Pseudescherichia, Pseudocitrobacter, Pseudomonas, Psychrobacter, Rahnella, Raoultella, Roseburia, Rosenbergiella, Rouxiella, Ruminiclostridium, Ruminococcus, Salmonella, Serratia, Siccibacter, Tatumella, Trabulsiella, Xenorhabdus* and *Yersinia*), suitable for visualisation in Archeopteryx [41]. The species definitions for these genomes were corrected and made strictly monophyletic to create the “NCBI_r88_Human17k” index.

